# Arp2/3 complex-dependent actin remodeling is required for efficient RSV uncoating in A549 cells

**DOI:** 10.64898/2026.03.11.711013

**Authors:** Enyu Xie, Maren Schubert, Jennifer Fricke, Benedikt Seligmann, Matthias Schaks, Mathias Müsken, Anika Steffen, Theresia E.B. Stradal, Sibylle Haid, Klemens Rottner, Christian Sieben

## Abstract

Productive human respiratory syncytial virus (RSV) cell-entry requires coordinated interactions between viral proteins and host-cell factors at the plasma membrane-actin cortex interface. Branched actin networks remodel this interface, but their precise contribution to the early stages of RSV infection remains unclear. Here, we interfered with Arp2/3 complex-dependent actin filament branching by generating A549 cell lines disrupted for expression of the essential Arp2 subunit by CRISPR/Cas9. Permanent loss of Arp2 reduced the infection of the RSV long GFP reporter virus as quantified over the first 24 h post-infection. Compromised infection efficiency in Arp2 knockout cells persisted at later time points and also resulted in reduced syncytia formation. Notably, these infection phenotypes were not accompanied by obvious changes in viral host cell attachment. Moreover, photoactivated localization microscopy (PALM) studies revealed comparable receptor diffusion and clustering in cells stably expressing mEos3.2-tagged insulin-like growth factor I receptor (IGF1R). Although Arp2/3-deficient cells displayed fewer albeit larger macropinosomes as compared to WT cells, no changes were observed for internalized RSV genome levels. In contrast, Arp2-deficient cells appeared suppressed in viral uncoating efficiency. Consequently, viral mRNA expression and the cellular type III interferon response were reduced. Together, these data reveal that Arp2/3 complex-dependent, branched actin networks contribute to the efficiency of RSV uncoating.

**Importance:** Human respiratory syncytial virus (RSV) is a major cause of severe respiratory disease. The infection initiates at the plasma membrane-actin cortex interface, yet the role of actin in productive RSV entry has remained unclear. Using CRISPR/Cas9 disruption of the essential Arp2/3 complex subunit Arp2 in A549 cells, we show that branched actin networks are required for efficient RSV infection. Despite actin network remodeling, photoactivated localization microscopy showed unchanged diffusion and clustering of the RSV receptor IGF1R. Although macropinocytosis was affected in Arp2-deficient cells, RSV attachment and internalization were not influenced. In contrast, a β-lactamase virus-like-particle-based assay revealed a defect in uncoating, followed by reduced viral gene expression and a weaker type III interferon response. These findings define Arp2/3 complex-dependent branched actin networks as a host determinant of RSV uncoating and provide a practical approach to quantify uncoating without engineering the RSV genome.

## 1 Introduction

Human respiratory syncytial virus (RSV) is an enveloped, negative-sense RNA virus in the genus *Orthopneumovirus* (family *Pneumoviridae*) [1]. It is a leading cause of acute lower respiratory tract infection. Its infection is initiated by attachment to the host cell plasma membrane. RSV uses its fusion (F) protein and attachment glycoprotein (G) to engage with the cell surface [2, 3], although G is dispensable for establishing infection in monolayer cell culture models [4]. As an obligate intracellular pathogen, RSV must transfer its ribonucleoprotein complex into the cytosol for replication. The delivery is achieved by F-mediated fusion of viral and host membranes, which can occur either at the plasma [5, 6] or the endosomal membrane following macropinocytic uptake [7]. Several host proteins have been proposed to function as receptor proteins in RSV entry, including IGF1R [8] and nucleolin [9]. Actin polymerization has been reported to promote nucleolin presentation at the cell surface [10]. Disruption of actin dynamics impairs RSV infection [6, 7]. However, the key actin-dependent mechanisms that enable productive virus entry and propagation have remained unclear.

Actin is one of the most abundant cytoskeletal proteins in eukaryotic cells. It polymerizes into microfilaments that provide structural support and enable dynamic processes. Actin assembly is most prominently driven by two classes of actin assembly factors: Formins, elongating individual or bundles of linear filaments and Arp2/3 complex best known for its generation of branched actin filaments [11, 12]. Branched actin networks are essential for sheet-like plasma membrane protrusions and contribute to the cells’ cortex, a thin peripheral layer beneath the plasma membrane [13, 14]. The actin cortex supports membrane remodeling events that can influence endocytosis [15], membrane tension [16], and surface-protein organization [17, 18]. In cells, the plasma membrane and cortex operate as a coupled mechanical signaling system. They act as a physical barrier for incoming viruses including RSV, which was shown to also exploit them to promote entry and cytosolic access [19]. However, the specific role of branched actin filaments in the earliest productive steps of RSV infection remains unclear. Aside from plasma membrane remodeling, Arp2/3 complex-driven actin networks are also implicated in vesicle and endomembrane trafficking[20], but how such events contribute to specific intracellular pathogen interactions with their hosts is also ill-defined.

In this study, we generated Arp2/3-deficient A549 cells to define how branched actin networks contribute to RSV infection at early stages. We disrupted the essential Arp2/3 complex subunit Arp2 using CRISPR/Cas9 in A549 lung epithelial cells. We showed that loss of Arp2 reduced RSV infection at early stages and limited syncytia formation at later time points. Remarkably, virus binding, receptor availability and viral internalization were unchanged. In contrast, a β-lactamase reporter assay revealed impaired uncoating in Arp2-deficient cells. Consequently, downstream viral transcription and type III interferon induction were also reduced. Together, these data identify Arp2/3 complex-dependent actin branching as crucial for efficient RSV uncoating.

## 2 Results

### 2.1 Arp2 knockout impedes RSV infection at both early and late stages

To investigate the role of the Arp2/3 complex in human respiratory syncytial virus (RSV) infection, Arp2/3 complex-deficient A549 cell lines were generated. The Arp2/3 complex comprises seven subunits: two actin-related proteins (Arp2 and Arp3) and five accessory subunits [21]. Arp2 and Arp3 form the actin-like catalytic core that nucleates branched filaments [22]. All seven subunits are required to maintain Arp2/3 complex integrity and support its proper function [23–25]. Therefore, the single *ACTR2* gene (encoding Arp2; NCBI ID 10097) was disrupted by CRISPR/Cas9 to permanently ablate Arp2/3 complex function (**Fig. S1A-C**). Five single-cell Arp2 knockout (KO) clones were initially employed for infection experiments.

These KO cell clones were challenged with RSV long strain expressing GFP (RSV-GFP) [26]. Initial screening for infection efficiency using image-based quantification revealed significantly impaired first-round infection across all KO clones **(Fig 1A and S1D)**. For more detailed experiments, two clones, Arp2 KO16 and KO17 were chosen. Flow cytometry showed approximately 50% fewer RSV-positive cells in Arp2 KO clones than WT at both 8 h and 12 h post-infection (p.i.) **(Fig 1B)**. At 48 h p.i., the fraction of RSV-GFP-positive cells decreased by approximately 40% in Arp2 KO cells compared to WT (**Fig. 1C-D**). In addition, syncytia were less frequent and smaller in Arp2 KO cells: WT, 27.2% of infected cells formed syncytia with a mean of 6.2 nuclei per syncytium; KO16, 9.8% (3.7 nuclei per syncytium); KO17, 6 % (4.3 nuclei per syncytium). These findings indicated that Arp2 KO impedes RSV infection mainly at early stages, thereby constraining downstream infection propagation.

**Fig. 1.**
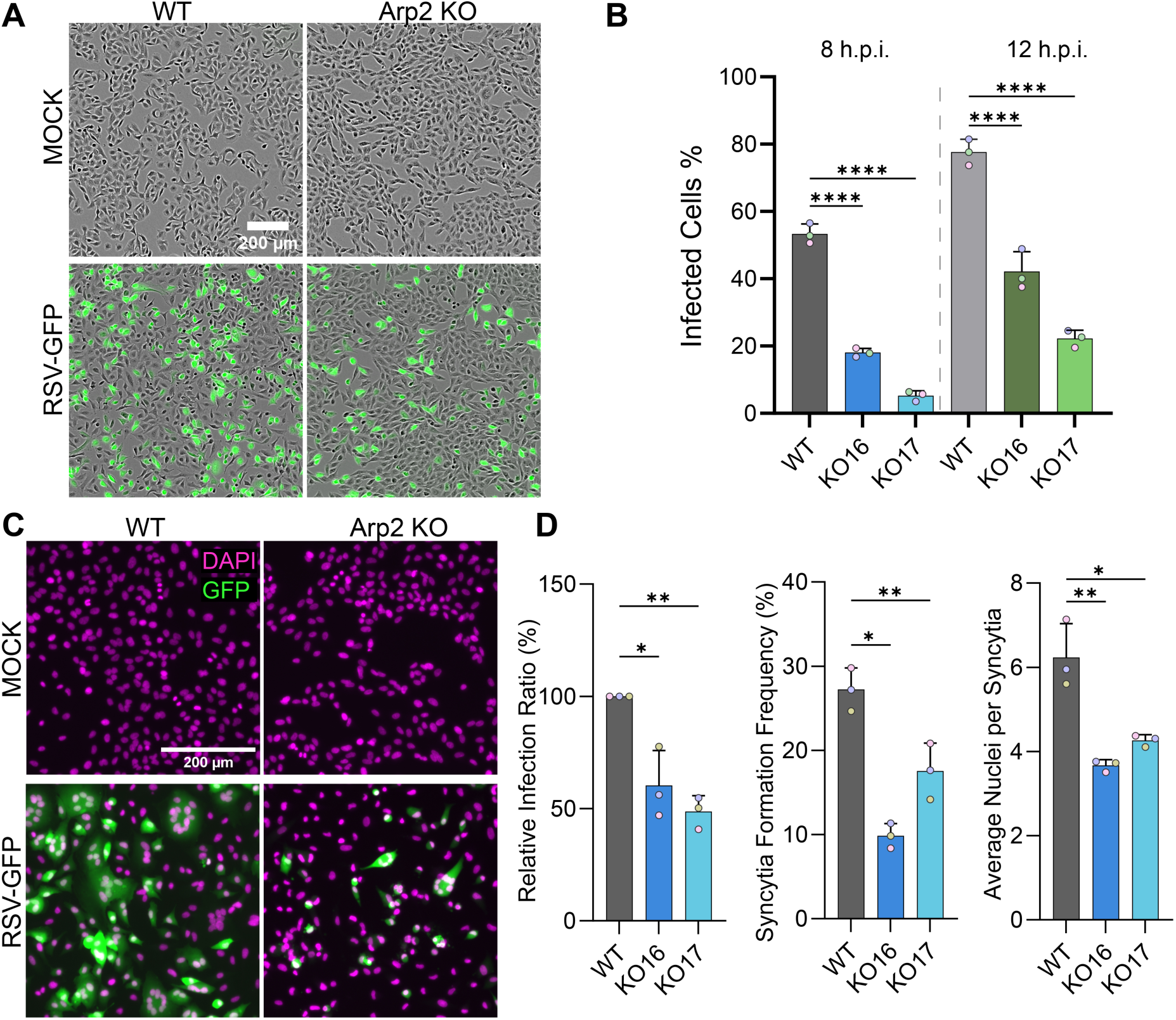
Arp2 KO impedes RSV infection in A549 cells at both early and late stages. (A) Representative images of RSV-GFP (MOI = 3) infected or mock-infected A549 WT and Arp2 KO17 cells at 24 h p.i.. Images were acquired on an Incucyte automated microscope using a 4 × objective. Infected cells were green due to GFP expression. Merged phase-contrast and GFP images are shown. Scale bar, 200 µm. (B) Flow cytometry analysis of RSV-GFP infection at 8 and 12 h p.i. in A549 WT cells and two Arp2 KO cell clones (MOI = 4). Data from three independent experiments (Mean ± SD; n = 3 biological replicates). Each dot represents the mean value of one experiment. ****p < 0.0001 (Two-way ANOVA). (C) Representative fluorescence images of RSV-GFP (MOI = 0.2) infected and mock-infected WT and Arp2 KO16 cells at 48 h p.i.. Cells were fixed and stained with Hoechst 33342 (nuclei; in pseudocolor magenta) before imaging. Merged images of DAPI and GFP are shown. Scale bar, 200 µm. (D) Quantification of infection and syncytium formation from the image series in (C) through CellProfiler. Infection was expressed as the percentage of GFP-positive cells and normalized per experiment to WT. A GFP-positive cell containing ≥3 nuclei was considered as syncytium. Syncytia formation frequency was calculated as the number of syncytia divided by the number of total infected cells. Data from three independent experiments (Mean ± SD; n = 3 biological replicates). Each dot represents the mean value of one experiment. *p < 0.05, **p < 0.01 (Two-way ANOVA).

### 2.2 Arp2 knockout does not influence RSV-cell attachment or local receptor dynamics

Since our data indicated a defect early in infection in Arp2 KO cells, we set out to pinpoint the specific infection stage that was impeded. RSV infection is initiated with virus attachment to cells and then engages entry factors, including insulin-like growth factor 1 receptor (IGF1R) and a co-receptor, nucleolin [8, 9]. Diminishing the branched actin nucleator Arp2/3 complex was shown to change cell morphology and physiology [27, 28], including endocytosis [13] and cell surface distribution of receptors [17, 18]. We therefore first examined RSV attachment and receptor mobility at the cell surface of WT versus Arp2-deficient cells.

To test whether loss of Arp2/3 complex activity limits RSV-attachment, we incubated cells with RSV-GFP on ice (MOI = 1) to permit binding while blocking entry. Image-based quantification across hundreds of cells showed no difference in the amount of bound virions per cell between Arp2 KO and WT cells (**Fig. 2A-B**). These data indicated that virus attachment is not influenced in Arp2 KO cells. Next, we took a closer look at receptor expression and organization. Using immunoblot analysis, we determined similar IGF1R and nucleolin protein levels in all cell types (**Fig. 2C**). Next, we used TIRF-PALM (total internal reflection microscopy-photoactivated localization microscopy) to determine the mobility of the RSV-receptor IGF1R tagged to mEos3.2 on the basal membrane of WT and Arp2 KO-cells (**Fig. 2D and S2**). Single particle tracking photoactivated localization microscopy (sptPALM) revealed comparable IGF1R diffusion rates (WT, 0.16 ± 0.05 µm^2^/s; KO16, 0.18 ± 0.06 µm^2^/s; KO17, 0.18 ± 0.05 µm^2^/s; **Fig. 2E-F**). Furthermore, PALM data of fixed cells revealed no detectable difference in IGF1R cluster size (**Fig. 2G-H**). Taken together, despite significantly reducing infection rates, Arp2 KO did not impair virus attachment or IGF1R dynamics at the plasma membrane.

**Fig. 2.**
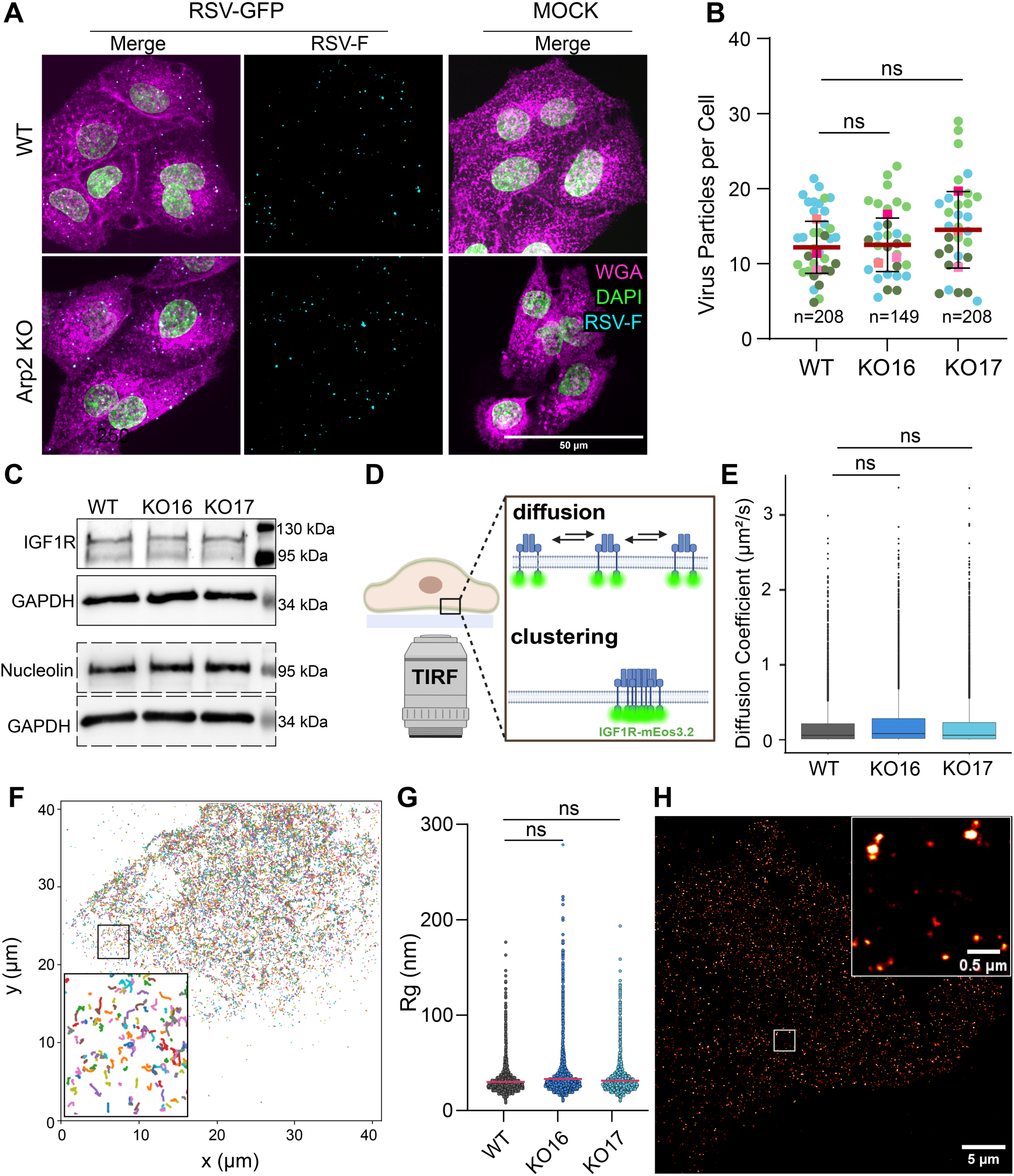
RSV attachment and cell receptor dynamics are unaffected in Arp2 KO cells. (A) Representative confocal microscopy images of RSV-GFP (MOI = 1) bound to A549 WT and Arp2 KO17 cells. Cells were incubated with virus or mock-infected for 1 h on ice, washed to remove unbound virus, fixed, and stained with Palivizumab (anti-RSV-fusion protein, in pseudocolor cyan), Hoechst 33342 (cell nuclei, in pseudocolor green), and WGA (wheat germ agglutinin-Alexa Flour 647, magenta). Images were acquired on a spinning-disk microscope and are shown as maximum-intensity projections of respective channels. Scale bar, 50 µm. (B) Quantification of viral binding from the image series in (A). Pooled data of three individual experiments (three biological replicates per experiment). n denotes the number of cells analysed. Experiments are color-coded; each dot represents one image, and each rectangle represents the experiment mean. Mean ± SD. ns, not significant (Two-way ANOVA). (C) Immunoblot of endogenous IGF1R and nucleolin in A549 WT and Arp2 KO cells. GAPDH (glyceraldehyde-3-phosphate dehydrogenase) served as loading control. (D) Schematic of using PALM in combination with TIRF to assess IGF1R diffusion and clustering. This image was created using Biorender (License number: CT29AQJ52X). (E) Diffusion coefficient of A549 WT and Arp2 KO cells stably expressing IGF1R-mEos3.2. The data were obtained from sptPALM imaging. ns, not significant (unpaired two-tailed Student’s t-test). (F) Representative sptPALM reconstruction with tracks of individual IGF1R-mEos3.2 particles generated from 10,000 frames (frame interval 30 ms). Trajectories are color-coded. (G) Comparison of the IGF1R cluster size of A549 WT and Arp2 KO cells stably expressing IGF1R-mEos3.2. ns, not significant (unpaired two-tailed Student’s t-test). (H) Representative PALM image showing the surface distribution of IGF1R-mEos3.2 in an A549 WT cell.

### 2.3 Arp2 KO affects macropinocytic dextran uptake but not RSV internalization

As viral attachment and receptor dynamics were unchanged in Arp2 KO cells, we asked whether cell entry was affected. RSV entry can be accomplished through different routes, such as plasma membrane fusion or macropinocytosis [7], with the primary mode of RSV entry remaining unclear. As actin rearrangements are required for macropinocytosis [29], we next quantified macropinocytic uptake of 70 kDa fluorescein-conjugated dextran [30]. As shown by microscopy (**Fig. 3A**), Arp2 KO cells formed larger dextran-positive vesicles than WT cells (median area 0.162 µm^2^ vs 0.108 µm^2^), yet displayed approximately 43% fewer vesicles per cell (**Fig. 3B**). As a result, the per-cell integrated dextran signal was modestly increased in Arp2 KO cells.

**Fig. 3.**
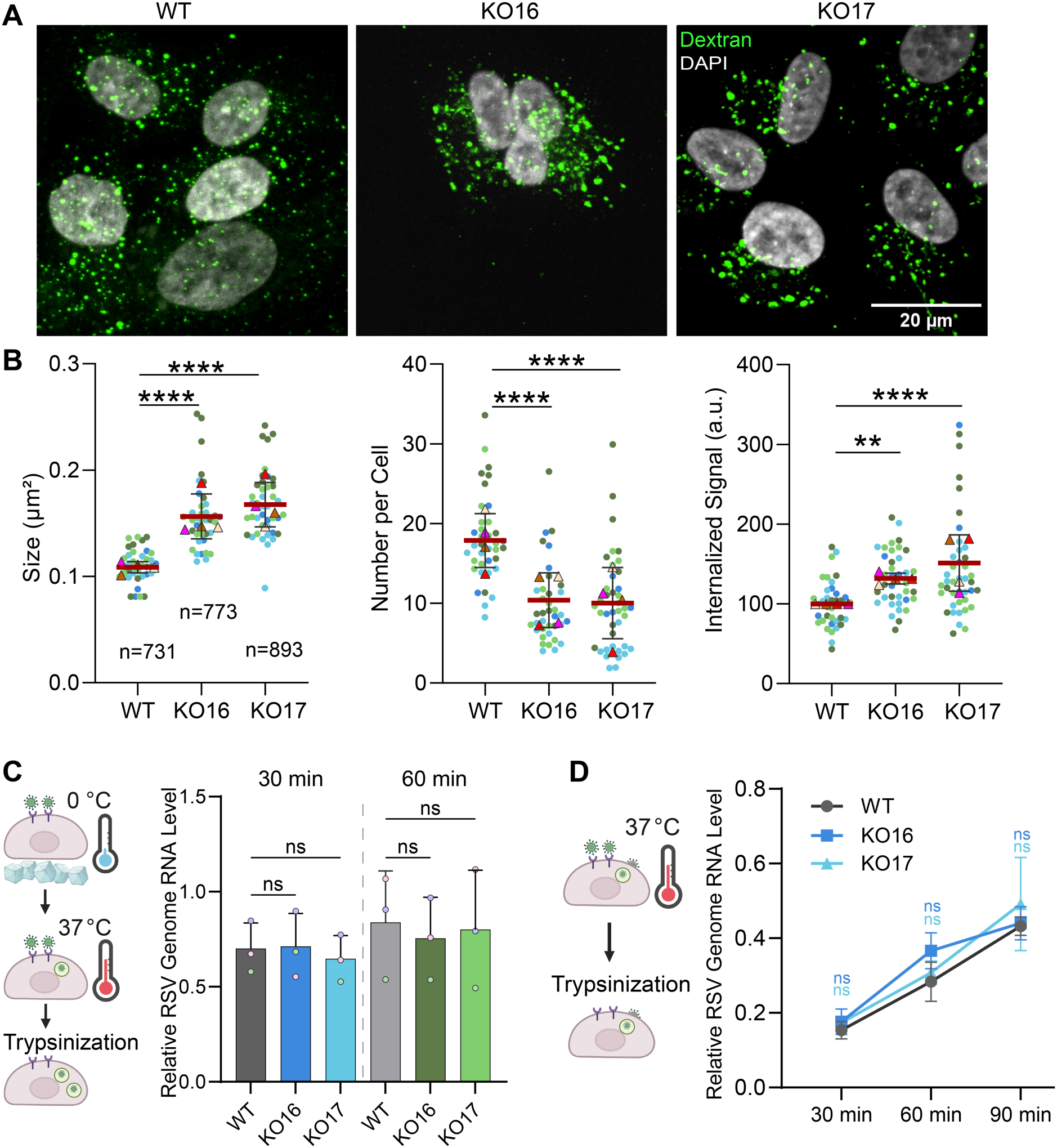
Arp2 KO affects macropinocytic dextran uptake but not RSV internalization. (A) Representative confocal microscopy images showing uptake of 70 kDa fluorescein-labelled dextran in A549 WT and Arp2 KO cells. Cells were incubated with 0.6 mg/ml dextran for 15 min. Cells were fixed and stained with Hoechst 33342 (shown in pseudocolor grey). Images were acquired on a spinning-disk confocal microscope and are shown as maximum-intensity projections of merged channels. Scale bar, 20 µm. (B) Quantification of dextran uptake from the image series in (A). Data are from four independent experiments (three biological replicates per experiment). n denotes the number of cells analysed. Experiments are color-coded; each dot represents one image, and each triangle indicates the experiment mean. Bars show mean ± SD across experiments. ns, ** P < 0.01; **** P < 0.0001 (Two-way ANOVA). a.u., arbitrary units. (C) RT-qPCR results of entry-assay with on ice preincubation. Cells were incubated with RSV (MOI = 5) on ice for 1 h; unbound virus was removed by washing. Cells were shifted to 37 °C for 30 or 60 min, then treated with trypsin before RNA extraction and RT-qPCR. RSV nucleoprotein RNA was quantified relative to the cellular housekeeping gene *HPRT1* (hypoxanthine phosphoribosyltransferase 1) RNA levels. Combined data from three independent experiments (n = 3 biological replicates). Each color represents one experiment; each dot is the experiment’s mean. Mean ± SD. ns, not significant (Two-way ANOVA). The scheme was generated in BioRender (License number QY29DGQSJZ). (D) RT-qPCR results of entry assay without on ice preincubation. Cells were incubated with RSV (MOI = 1) at 37 °C for 30, 60 or 90 min. The non-internalized virus was removed by trypsin. Cells were then collected for RNA extraction and RT-qPCR. RSV nucleoprotein RNA was quantified relative to *HPRT1*. Representative data from two independent experiments (n = 3 biological replicates). Mean ± SD. ns, not significant (One-way ANOVA). Dark-blue denotes statistics for WT vs KO16; light-blue denotes statistics for WT vs KO17. The scheme was generated in BioRender (License number QY29DGQSJZ).

We next tested whether these changes affect RSV internalization. After on-ice binding, cells were warmed to 37 °C for virus internalization. At 30 or 60 min p.i., non-internalized virions were removed by trypsin, and intracellular viral RNA was measured by RT-qPCR. We found that Arp2-KO and WT internalized similar amounts of virus (**Fig. 3C**). Because infection without on-ice pre-incubation can favour direct fusion in NHBE cells [6], we repeated the experiment without the on-ice binding step. The latter conditions also confirmed an increased internalization of virus over time, but again with no difference between Arp2 KO and WT cells (**Fig. 3D**). Together, Arp2 loss altered macropinocytic uptake, but did not affect virus entry.

### 2.4 Arp2 knockout impedes RSV uncoating and downstream transcription

Next, we asked whether reduced infection in Arp2 KO cells stems from impeded virus uncoating. To measure uncoating, we used β-lactamase-containing virus-like particles (VLPs). This approach was adapted from β-lactamase (BlaM)-based entry assays originally developed for HIV-1 (human immunodeficiency virus type 1) [31] and SARS-CoV-2 (severe acute respiratory syndrome coronavirus 2) [32]. SARS-CoV-2 membrane protein (M) and SARS-CoV-2 envelope protein (E) were used for assembly of the VLPs. As the RSV fusion protein (F) alone is sufficient to phenocopy RSV entry patterns in cell culture [33], we produced VLPs displaying RSV F on the surface (**Fig. 4A**). BlaM was fused to the SARS-CoV-2 nucleocapsid protein (N) to enable incorporation into VLPs. Transmission electron microscopy showed that the VLPs are morphologically heterogeneous **(Fig. S3A)**. Additionally, immunoblotting confirmed the incorporation of RSV F and BlaM-N **(Fig. S3B).** Upon fusion, VLPs uncoat and release BlaM into the cytosol, where it cleaves the FRET (Förster resonance energy transfer)-based substrate CCF2-AM added separately during the experiment. This generates a signal quantifiable by flow cytometry [34] (**Fig. 4A**). Interestingly, Arp2 KO cells showed a strong reduction in BlaM-positive cells following VLPs infection, with clonal variability but an average decrease to approximately 50% of WT (**Fig. 4B**).

**Fig. 4.**
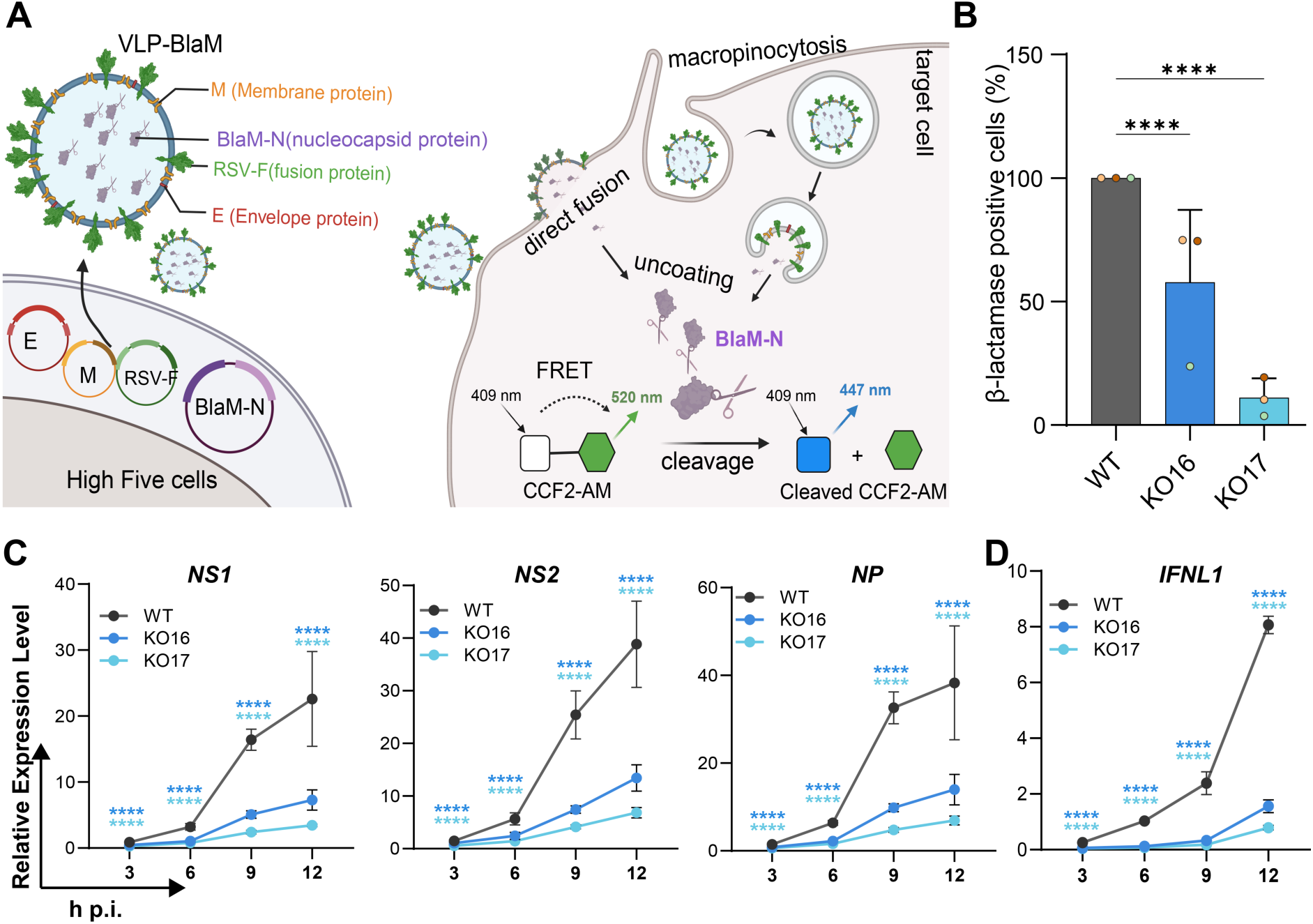
Arp2 knockout impedes RSV uncoating and downstream transcription. (A) Schematic of using VLP-BlaM to assess RSV uncoating. VLPs were generated using a SARS-CoV-2-based VLP system [32] bearing the RSV fusion protein instead of Spike, and was applied to A549 WT or Arp2 KO cells. Cytosolic BlaM activity was assessed using CCF2-AM, a FRET-based substance. This image was created using Biorender (License number HC29AQJKBO). (B) Arp2 KO impeded RSV uncoating. Cells were infected with VLPs or mock infected at 37 °C for 2 h, then incubated with CCF2-AM in the dark at RT for 3 h. Cells were detached, fixed in PFA, and analyzed using flow cytometry. The percentage of BlaM-positive cells was normalized per experiment to A549 WT. Data are from three independent experiments (n = 3 biological replications). Mean ± SD. **** P < 0.0001 (Two-way ANOVA with Dunnett’s multiple comparisons test) (C) Arp2 KO reduced RSV transcription. RT-qPCR time course of RSV gene expression following infection (RSV-GFP, MOI = 0.5) in A549 WT versus Arp2 KO cells. RSV *NS1* (non-structure protein 1), *NS2* (non-structure protein 2), *N* (nucleoprotein) transcripts were quantified and normalized to cellular housekeeping gene *HPRT1*. Representative data from two independent experiments (n = 3 biological replicates). Mean ± SD. ****p < 0.0001 (one-way ANOVA). Dark-blue asterisks denote statistics for WT vs KO16; light-blue asterisks denote statistics for WT vs KO17. (D) RSV induced a milder type III interferon response in Arp2 KO cells compared to WT cells. RT-qPCR time course of *IFNL1* expression following RSV infection (MOI = 0.5). The transcripts were quantified and normalized to *HPRT1*. Mean ± SD. ****p < 0.0001 (one-way ANOVA). Dark-blue asterisks denote statistics for WT vs KO16; light-blue asterisks denote statistics for WT vs KO17.

Since we found that virus uncoating was affected in Arp2 KO cells, we sought to test next if the cells also display reduced expression of viral proteins as a consequence of fewer viral genomes being released. To this end, cells were infected with RSV-GFP (MOI = 0.5), and the transcription levels of viral and cellular genes were analysed by RT-qPCR. Notably, we detected markedly lower viral mRNA levels in Arp2 KO cells than in WT cells (**Fig. 4C**). Finally, we monitored innate immune responses. Consistent with a previous report [35], type III interferon was the dominant interferon induced by RSV infection, whereas type I interferon remained very low throughout all cell types (**Fig. S3D**). Importantly, a weaker type III interferon response was observed in Arp2 KO cells (**Fig. 4D**). Taken together, these data suggest a defect in RSV uncoating in Arp2 KO cells that consequently also limited viral transcription and induction of the type III interferon response.

## 3 Discussion

Actin remodeling is widely implicated at the early stages of virus infection [7, 10]. Branched actin networks nucleated by the Arp2/3 complex are considered as key drivers of cortical architecture and plasma membrane remodeling [36, 37]. However, the contribution of branched actin to RSV infection has remained unclear. In this study, we identified Arp2/3 complex-dependent branched actin as a determinant of efficient RSV uncoating in A549 cells.

Here, CRISPR/Cas9-edited Arp2 knockout (KO) A549 cells were used to study the role of Arp2/3 complex during RSV infection. We found reduced RSV infection during the first 24 h post-infection in Arp2 KO cells **(Fig. 1A-B; Fig. S1D)**. This defect was reproducible across independent experiments. However, prior work has reported mixed effects of Arp2/3 complex perturbation on RSV infection. In HeLa cells, CK-869 (an Arp2/3 complex inhibitor and analogue of CK-666) reduced RSV internalization and infection at 6 h p.i. (around 40%), consistent with our findings [7]. In contrast, Kieser *et al*. reported that CK-666 (30 µM, 4 h) added into the inoculum did not affect the first-round of RSV infection in 1HAEo-cells [10]. In addition, Mehidi *et al.* reported that *Arp2* siRNA in A549 cells had little effect on entry or early viral gene expression, but impeded cell-to-cell spread at later stages [38]. Several factors may explain the differences to our results. First, the effects of distinct inhibitors strongly depend on dose and exposure time. Second, genetic knockout typically produces a more complete and sustained loss of function than siRNA [39].

Arp2/3 complex inhibition was previously described to induce blebbing (at the expense of lamellipodia formation and membrane ruffling) in multiple cell types [40–43]. Such remodeling may well perturb the spatial organization and lateral mobility of plasma membrane proteins [17, 18]. However, these morphological changes did not alter RSV attachment (**Fig. 2A-B**). We also observed similar IGF1R dynamics at the basal membrane by super-resolution microscopy (**Fig. 2E-H**), suggesting comparable availability of this entry factor in cells with and without Arp2/3 complex activity. One limitation of our study could be that we did not directly assess RSV-induced receptor rearrangements, including IGF1R clustering or nucleolin recruitment at the virus-cell contact site. Nucleolin is a co-receptor that translocates from the nucleus to the plasma membrane and supports RSV infection [8]. Actin remodeling is required for nucleolin translocation [10]. However, the observed unchanged viral internalization supports a preserved functional receptor availability in Arp2 KO cells at the plasma membrane (**Fig. 3C-D**).

The Arp2/3 complex has been reported to have variable effects on macropinocytosis. It was shown to be recruited to macropinocytic cup formation sites within seconds [44]. Moreover, inhibition of Arp2/3 complex can reduce macropinosome formation and delay vacuolar rupture [45]. In contrast, other studies reported that CK-666 increased dextran uptake intensity [46] or had little effect on macropinocytosis [38]. In our study, Arp2 KO produced fewer but larger dextran-positive macropinosomes (**Fig. 3A-B**). A modest increase in integrated uptake was also observed. This altered pattern may reflect delayed cup closure or scission, allowing more cargo to accumulate per vesicle. Despite these changes, our RT-qPCR-based entry assay showed unperturbed RSV internalization (**Fig. 3C-D**). This assay captured the net outcome of internalization but did not distinguish the entry route. Thus, the result could be explained in several ways. First, RSV may require only a small fraction of the available macropinocytic machinery. Therefore, moderate changes in vesicle number and size do not limit particle internalization. Second, the increased vesicle size could partially compensate for the reduced vesicle number, resulting in similar net capacity for RSV internalization. Third, RSV can enter cells via plasma membrane fusion or endocytic uptake; these parallel routes may buffer perturbations to either pathway.

In this study, we applied β-lactamase carrying VLPs to quantify RSV uncoating (**Fig. 4A**). These particles can be generated rapidly by co-transfecting a small plasmid set. They are replication-incompetent, which enhances biosafety and reduces experimental complexity. A limitation of the assay is that it reports total cytosolic delivery. It is unable to distinguish direct fusion at the plasma membrane and indirect fusion at the endosome membrane. In addition, it measures the uncoating efficiency of VLPs rather than authentic RSV. Nevertheless, this assay revealed reduced uncoating in Arp2 KO cells (**Fig. 4B**). This defect aligned with reduced viral gene expression (**Fig. 4C**) and a weaker type III interferon response (**Fig. 4D**) after infection with authentic RSV.

Several mechanisms could explain the requirement for Arp2/3 complex-dependent actin filament branching during RSV uncoating. First, branched actin may support fusion pore expansion by providing mechanical support at the cortex. Consistent with this model, Arp2 KO cells frequently exhibit blebbing, which is associated with weakened cortical networks and reduced membrane-cortex coupling [41, 47]. In this scenario, fusion may initiate, but fusion pore expansion and ribonucleoprotein release would be inefficient. Second, loss of branched actin could delay the trafficking of host factors to macropinosomes, impairing their maturation and, in turn, restricting RSV uncoating. Consistent with this idea, macropinocytosis appeared modified in Arp2 KO cells. Moreover, delayed macropinosome rupture was reported after Arp2/3 complex inhibition [45]. Further work is needed to pinpoint the branched actin-dependent step and to resolve the relative contributions of fusion at the plasma versus endosomal membrane.

## 4 Methods

### Cell lines and culture conditions

Human lung epithelial A549 cells (ATCC, CCL-185), HEp-2 cells (ATCC, CCL-23), and HEK-293T cells (human embryonic kidney 293T; DSMZ, ACC 635) were maintained in Dulbecco’s modified Eagle’s medium (DMEM; Gibco, 61965026) supplemented with 10% heat-inactivated fetal bovine serum (Sigma-Aldrich, F7524-500mL). For the RSV infection assay, cells were incubated with infection medium (DMEM + 2% FBS) during and after infection. All mammalian cells were grown at 37 °C in a humidified incubator with 5% CO_2_. High Five insect cells (BTI-TN-5B1-4) were cultured with EX-CELL 405 medium (Merck, 24405C), and kept at a cell density of 0.3-5.5 × 10^6^ / mL (27 °C, 150 rpm, without CO_2_).

### Antibodies

The antibodies used in this study were as follows: rabbit recombinant monoclonal anti-Arp2 antibody (1:2,000; EPR7979, ab129018, Abcam); mouse monoclonal anti-HSC70 antibody (1:5,000; sc-7298, Santa Cruz); GAPDH (D16H11) rabbit monoclonal antibody (1:1,000, 5174S, Cell Signaling Technology); IGF1R polyclonal antibody (1:500, BS-0227R, Bioss); nucleolin monoclonal antibody (1:500, 39-6400, Invitrogen); anti-Beta lactamase antibody (1:500, ab12251, Abcam); anti-SARS-CoV-2 nucleocapsid protein antibody (1:500, ABK79-E11-M, Abcalis); Palivizumab (10 µg/mL, a gift from Dr. Thomas Pietschmann, TWINCORE, Germany); HRP-conjugated goat anti-rabbit antibody (1:10,000; G-21234, Invitrogen); HRP-conjugated goat anti-mouse antibody (1:10,000; G-21040, Invitrogen); goat anti-Human cross-adsorbed secondary antibody, Alexa Fluor™ 488 (1:200, A-11013, Invitrogen); goat anti-human IgG cross-adsorbed secondary antibody, HRP (1:10,000; A18811, Invitrogen).

### Generation of A549 Arp2 (*ACTR2*) CRISPR/Cas9 knockout cells

PX459-Arp2 was constructed by cloning a sgRNA (single-guide RNA) targeting *ACTR2* (5’-TCATTCCAGTTTGTGAAGTG-3’) into pSpCas9-(BB)-2A-Puro plasmid v2.0 (PX459; Addgene, 62988) (**Fig. S1A**), as described [48]. A549 cells were transfected with PX459-Arp2 using jetOPTIMUS (101000006, Sartorius / Polyplus) according to the manufacturer’s protocol. At 48 h post-transfection, cells were transferred into selection medium containing 1.5 µg/mL puromycin for 96 h. Subsequently, surviving cells were diluted for single-cell colony formation into conditioned medium (30%) containing Rho-Kinase inhibitor (Y27632, 10 µM), the latter of which is known to counteract anoikis. Single-cell colonies were further expanded and explored for the absence of Arp2 expression by immunoblotting **(Fig. S1B)**. Clones identified to lack Arp2 expression by Western Blotting were expanded further and subjected to genomic DNA extraction followed by PCR-amplification of the target locus and TIDE-sequencing using specified sequencing primer, to confirm the absence of a WT allele or presence of any in-frame deletions or insertions (see **Table S1 and Figure S1C**).

### Western blot analysis

Cells were lysed and boiled at 95 °C for 5 min. Proteins were separated by 10% SDS-PAGE and transferred to nitrocellulose membranes. Membranes were blocked in 5% skimmed milk for 1 h and then probed with primary antibodies. HPR-conjugated secondary antibodies were then applied (1:10,000), respectively. Bands were visualized using SignalFire™ ECL Reagent (Cell Signaling Technology, 6883) and imaged by ECL Chemostar (Intas).

### Stable expression of IGF1R-mEos 3.2 by lentiviral transduction

pGenLenti-IGF1R-mEos3.2 construct contains a codon-optimized human IGF1R (Insulin-like growth factor 1 receptor, Genbank AB463073.1). IGF1R was fused C-terminally to mEos3.2 [49] via the linker sequence GGGPVPQWEGFAALLATPVAT [50]. The construct was custom-synthesized and cloned into the pGenLenti backbone by Genscript. pCAGGS-VSV-G (a gift from Dr. Roland Schwarzer, University Hospital Essen, Germany), pCMV-R8.74 (Addgene, 22036) and pAdvantage (Promega, E1711) were used for lentiviral packaging. In short, HEK-293T cells were seeded into T25 flasks and transfected at 60-70% confluence with 2.3 µg pCMVR8.74, 2.3 µg pGenLenti-IGF1R-mEos3.2, 0.23 µg pCAGGS-VSV-G, 0.5 µg pAdvantage, and 21 µg linear polyethylenimine (PEI, Sigma-Aldrich, 919012-100MG). At 16 h p.t. (post-transfection), sodium butyrate (final concentration 500 mM; Thermo Fisher) was added for 3 h, after which the medium was refreshed. At 32 h p.t., the supernatant was collected, clarified by a 0.45 µm syringe filter, and applied to A549 WT and Arp2 KO cells. Transduced cells were selected with 1 µg/mL puromycin for 7 days.

### RSV propagation and purification

RSV long GFP strain (RSV-GFP) was a gift from Dr. Marie-Anne Rameix Welti (Institut Pasteur, France)[26]. Confluent HEp-2 monolayers were infected by RSV-GFP at MOI = 0.01-0.1 for 2 h, and then maintained in infection medium for 48-72 h. When 70-90% cells showed cytopathic effects, cells and supernatant were harvested, vortexed for 2 min, and clarified through centrifugation (1000 × g, 10 min, 4 °C). The supernatant was then mixed with 10% (v/v) cryo-conservation solution (HEPES 0.5 M, MgSO₄ 1 M, pH 7.5) and snap-frozen in an ethanol/dry-ice bath before storage at −80 °C.

For purification, the virus stock was overlaid on a discontinuous sucrose gradient (30%, 45%, 60% sucrose in NTE buffer), as described previously [51]. After centrifugation (150,000 × g, 1.5 h, 4 °C; SorvallTM WX Ultra 80, Thermo Fisher), the RSV band (between 30% and 45% sucrose) was collected, aliquoted, snap-frozen, and stored at −80 °C.

RSV-GFP titers were determined by infecting confluent A549 WT cell monolayers in 96-well plates with serial dilutions. GFP-positive cells were quantified by Incucyte (Sartorius S3, Germany) at 24 h p.i. (hours post-infection). The titer was calculated as TU/mL (transducing units per milliliter) using the formula: (number of GFP-positive cells × fold of dilution) / the volume of virus added.

### Microscopy of RSV-GFP infected cells

6 × 10^4^ cells were seeded per well in 24-well plates one day before infection. Cells were incubated with RSV-GFP for 1 h (MOI = 3 for the 12 and 24 h p.i. dataset, **Fig. 1A and S1D**; MOI = 0.2 for the 48 h p.i. dataset, **Fig. 1C-D**), after which the inoculum was replaced by fresh infection medium. GFP-positive cells were imaged and quantified through Incucyte at 12 and 24 h p.i.. At 48 h p.i., cells were fixed, stained with Hoechst 33342 (0.1 µg/mL; Invitrogen, H21492), and imaged by EVOS (M5000, Thermo Fisher). The infection frequency was quantified in CellProfiler. A GFP-positive cell containing ≥3 nuclei was defined as a syncytium. Syncytia formation frequency was calculated as follows:

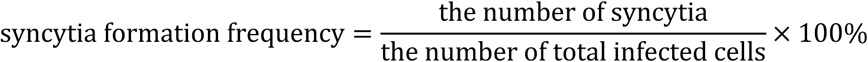

### Flow cytometry of RSV-GFP-infected cells

6 × 10^4^ cells were seeded per well in 24-well plates one day before infection. Cells were incubated with RSV-GFP (MOI = 4) in infection medium for 1 h, after which the inoculum was replaced by fresh infection medium. At 8 or 12 h p.i., cells were washed, trypsinized, and pelleted (300 × g, 5 min). Pellets were resuspended and fixed with 4% PFA/PBS (paraformaldehyde, 28908, Thermo Fisher Scientific) for 20 min at room temperature. Then cells were repelleted, washed, resuspended in FACS buffer (PBS + 2% FBS), and run through a BD LSR Fortessa (488 nm excitation; 525/50 nm emission). Gating: singlets by SSC-A /FSC-A and SSC-H; GFP threshold was set using mock-infected controls. Data were analyzed in FlowJo V10.

### Virus-cell binding assay

1 × 10^5^ cells were seeded onto 18 mm glass coverslips in 12-well plates one day before infection. Cells were washed three times with ice-cold PBS and incubated with RSV-GFP (MOI = 1) on ice for 1 h. After removing the inoculum, cells were washed three times with ice-cold PBS and fixed with 4% PFA. Fixed cells were permeabilized with 0.1% Triton X-100 (10 min) and blocked in 0.2% BSA (1 h, room temperature). Palivizumab was applied for 1 h, followed by goat anti-human IgG-Alexa Fluor 488. Nuclei were stained with Hoechst 33342 and cell membranes with wheat germ agglutinin-Alexa Fluor 647 (5 µg/mL; Invitrogen, W32466) for 15 min. Coverslips were mounted in ProlongTM Gold Antifade reagent. Cells were then imaged by a 100 × objective on a confocal spinning-disk microscope (CSU-W1, Nikon). Image analysis was performed in Fiji using particle analysis.

### Live-cell single-molecule imaging of IGF1R

Single-molecule tracking photoactivated localization microscopy (sptPALM) was used to probe insulin-like growth factor 1 receptor (IGF1R) diffusion, adapted from a published protocol [52]. Briefly, a sticky 8-well chamber (Ibidi, 80828) was mounted on a 3D epoxy-coated glass slide (PolyAn, 10400206). 25,000 cells stably expressing IGF1R-mEos3.2 were placed into each well, and cultured at 37 °C. Next day, cells were imaged at 37 °C in HEPES-buffered infection medium (without phenol red). Images were acquired on a Nikon Ti Eclipse N-STORM Microscope with an Apo TIRF 100x Oil DIC N2, NA 1.49 objective. To assess IGF1R-mEos3.2 localization, 10,000 frames were acquired at 30 ms intervals using a 405 nm laser for photoconversion and a 561 nm laser for excitation.

The locations of single emitters were detected using DECODE [53]. Sample drift was then corrected in MATLAB R2020b (The MathWorks Inc., Natick, MA). The first 10% frames and locations with certainty < 0.8 in either x or y directions were excluded. Trajectories were reconstructed in TrackPy [54] by linking remaining localizations using a KDTree algorithm (maximum search range = 0.8 pixels, zero-frame-gap memory). Trajectories lasting 0.3 - 4 seconds (5 - 133 frames) were included in the analysis. Diffusion coefficients were estimated by linear fitting in log space of the mean square displacement. Python packages NumPy [55], Matplotlib [56], pandas [57], plotnine [58] were used in Thonny (Python 3.10.11) for data visualization. Statistical analysis was performed using GraphPad Prism 10.

### Quantification of IGF1R nanoclustering

The protocol was adapted from a previous publication [59]. Briefly, 25,000 cells stably expressing IGF1R-mEos3.2 were seeded per well into a glass-bottom 8-well dish. On the day of imaging, cells were fixed with 4% PFA for 20 min and washed three times with PBS. Images were acquired on a Nikon Ti Eclipse N-STORM Microscope with an Apo TIRF 100x Oil DIC N2, NA 1.49 objective. 20,000 frames were acquired at 30 ms intervals using a 405 nm laser for photoconversion and a 561 nm laser for excitation.

The locations of single emitters were detected using DECODE [53]. Sample drift was then corrected using custom scripts in MATLAB R2020b (https://github.com/christian-7/SMLM_Tutorial). Localizations were clustered using the built-in DBSCAN function of MATLAB. Images were rendered using Thunderstorm v1.3 in Fiji. Statistical analysis and data visualization were performed using GraphPad Prism 10.

### Dextran uptake assay

Macropinocytosis was quantified by uptake of 70 kDa fluorescein-labelled, lysin-fixable dextran (0.6 mg/mL; Invitrogen, D1822), as described [30]. Briefly, cells were washed three times with PBS and then incubated with dextran (15 min, 37 °C). After washing three times with ice-cold PBS, cells were fixed, stained with Hoechst 33342, and wheat germ agglutinin-Alexa Fluor 647. Cells were then imaged with a 100 × objective on a confocal spinning-disk microscope (CSU-W1, Nikon). Image analysis was performed in Fiji using particle analysis.

### Virus entry assay

#### Entry assay with on-ice preincubation

The protocol was adapted from a previous publication [60]. Briefly, cells were seeded in 24-well plates one day before infection at a density of 6 × 10^4^ cells per well. The next day, cells were washed three times with ice-cold PBS and then challenged with RSV-GFP at MOI 5 on ice for 1 h. After binding, cells were washed three times with PBS and then incubated in infection medium at 37 °C for 0.5 or 1 h. Cells were then treated with ice-cold 0.25% trypsin-EDTA on ice for 10 min to remove non-internalized virions. Cells were then washed three times with PBS. The High Pure RNA Isolation Kit (11828665001, Roche) was applied to lyse cells and isolate RNA. The GoTaq® 1-Step RT-qPCR system (A6020, Promega) was used to quantify RSV nucleoprotein RNA. Cellular *HPRT1* mRNA served as a reference. Data were processed in LightCycler® 96 SW 1.1 (Roche Diagnostics). GraphPad Prism 10 was used for statistical analysis and data visualization.

#### Entry assay without preincubation

Cells were prepared as described above. On the day of the experiment, cells were washed once with infection medium and then challenged with RSV-GFP (MOI = 1) at 37 °C. After incubating for 30, 60 or 90 min, cells were washed with PBS. Trypsin-EDTA was applied to detach the cells and to remove the non-internalized virions from the cell surface. Cells were resuspended in growth medium and centrifuged at 300 × g for 5 min. The supernatant was aspirated and the pellet washed with infection medium and centrifuged again at 300 × g for 5 min. The cell pellets were then used for RNA isolation and RT-qPCR as described above.

### Generation of virus-like particles carrying β-lactamase

The virus-like particles carrying β-lactamase were produced from a SARS-CoV-2 (severe acute respiratory syndrome coronavirus 2) VLP production platform [32]. The constructs for VLP assembly (pOpiE2_SARS-CoV-2-E and pOpiE2_SARS-CoV-2-M) were prepared as described before [32]. pOpiE2_RSV-F was generated by subcloning the RSV fusion protein from pcDNA 3.1_RSV-F (a gift from Dr. Thomas Pietschmann, Twincore, Germany) [33] into a modified pOpiE2 backbone [61] containing the Spike packaging signal. pOpiE2_BlaM-SARS-CoV-2-N was constructed by fusing β-lactamase (BlaM) (from BlaM-Vpr; Addgene,21950) [31] via a short linker (GGGGGKGGR) to the N terminus of SARS-CoV-2 Nucleocapsid (Genebank UGC7399.1: 2-149aa, X16T, X40R), followed by its insertion into the pOpiE2 backbone. Transfection of High Five cells was performed as described previously [61]. In brief, pOpiE2_SARS-CoV-2-E, pOpiE2_SARS-CoV-2-M, pOpiE2_RSV-F, and pOpiE2_SARS-CoV-2-N-BlaM were mixed in equal amounts to get a plasmid mixture. For each 1 × 10^6^ cells, 1 µg plasmid mixture and 4 µg of 40 kDa linear PEI (1 mg/mL stock solution) were used. Plasmid mixture and linear PEI was added directly to the cells at a density of 4×10^6^ cells/mL. At 6-10 h p.t., fresh medium was added to adjust cell density to ∼1×10^6^cells/mL; at 48 h p.t., the culture volume was doubled. At 96 h p.t., supernatants were collected, and clarified through two centrifugation steps (180 x g, 4 min, RT followed by 2,000 × g, 10 min, 4 °C). VLPs were precipitated by mixing the supernatant with 0.136x volume of 50% (w/v) PEG6000 + 2.2% NaCl and incubating 2.5 h at 4 °C. Afterwards, the VLPs were pelleted at 4 °C, 20 min at 2000 x g and resuspended in NTE buffer [51]. To ensure a homogenous distribution, the resuspended VLPs were gently rotated at 18 °C for 1 h before 10% (v/v) cryo-conservation solution was added, and VLPs were snap-frozen and stored at −80 °C.

### Transmission electron microscopy of VLPs

VLPs were fixed in PFA (final concentration 6%) for 1 h at room temperature and further processed as previously described [62]. In brief, VLPs were adsorbed to a carbon film taken up with a copper grid, and washed twice in water before negative staining with 4% (w/v) aqueous uranyl acetate and drying with filter paper and on a light bulb. TEM was performed with a Zeiss Libra120 Plus transmission electron microscope (Carl Zeiss, Oberkochen, Germany) at an acceleration voltage of 120 kV at calibrated magnifications using ITEM Software (Olympus Soft Imaging Solutions, Münster).

### Uncoating assay

The uncoating assay was adapted from a published protocol [34]. Cells were seeded into 24-well plates 24 h before infection, with one extra well per condition reserved for cell counting. On the day of assay, the extra well was trypsinized to determine cell density. The inoculum was scaled to deliver an equivalent VLP-to-cell ratio across groups. Cells were then exposed to the same normalized amount of VLP-BlaM for 2 h at 37 °C. Subsequently, cells were washed with infection medium and then loaded with CCF2-AM (3 µM; Invitrogen, K1032) plus 2.5 mM Probenecid (AAB2001009, Thermo Scientific Chemicals), then incubated for 3 h in the dark at room temperature. Cells were washed three times with PBS, trypsinized, resuspended, pelleted (300 × g, 5 min), and fixed in 4% PFA for 20 min. Pellets were then washed and resuspended in PBS with 2% FBS for flow cytometry analysis. Data were processed using FlowJo™ v10.8 Software (BD, Ashland, OR). Statistical analysis and data visualization were performed using GraphPad Prism 10.

### RT-qPCR for detecting viral and cellular transcription levels

Cells were infected with RSV-GFP (MOI = 0.5) for 0, 3, 6, 9, and 12 h, respectively. At the indicated time point, cells were washed twice with PBS and detached by trypsin. Trypsinization was stopped by adding full growth medium, and the cell suspension was centrifuged at 300 × g for 5 min. The supernatant was discarded, and the cells were washed once with infection medium. Cells were re-pelleted through centrifugation, and then lysed using the High Pure RNA Isolation Kit. Total RNA was extracted as described above. cDNA synthesis and RT-qPCR were performed using the GoTaq® 2-Step RT-qPCR System (Promega, A6010) following the manufacturer’s instructions. Viral (*NS1, NS2, N*) and host (*IFNL1* and *IFNB*) transcripts were quantified with primers as provided **(Table S1)**; *HPRT1* served as the reference gene. Data were processed using LightCycler® 96 SW 1.1 (Roche Diagnostics). Statistical analysis and data visualization were performed using GraphPad Prism 10.

## Author Contributions

Study conception: E.X. and C.S.; Experiments: E.X., Mar.S., M.M., J.F., B.S., M.S., A.S., and C.S.; Data analysis and interpretation: E.X., C.S., K.R., S.H., T.E.B.S and M.M.; Manuscript drafting: E.X. and C.S.; Supervision and project administration: C.S.; All authors contributed to the scientific discussion and manuscript editing.

## Acknowledgements

The authors gratefully acknowledge Dr. Marie-Anne Rameix-Welti (Institut Pasteur, France) for providing RSV reporter viruses. We thank Dr. Thomas Pietschmann (Twincore, Germany) for providing reagents and for critically reading the manuscript. We also acknowledge Dr. Roland Schwarzer (University Hospital Essen, Germany) for providing plasmids and assistance with the uncoating assay. We are grateful to Dr. Cord Brakebusch (BRIC Copenhagen, Denmark) for valuable tips on single-cell cloning. We also thank Dr. Lothar Groebe (Flow Cytometry Platform, Helmholtz Centre for Infection Research, Germany) for technical assistance with flow cytometry, and Ina Brentrop (Microscopy Platform, Helmholtz Centre for Infection Research, Germany) for assistance with EM sample preparation. E.X. received a PhD fellowship from the China Scholarship Council (Grant No. 202106990020). C.S. acknowledges support by the Helmholtz Association (VH-NG-1526). K.R. and T.E.B.S. received funding from the German Research Council (Research Training Group GRK2223 and individual grant RO2414/8-1) and the Helmholtz Society, respectively.

