## Supplementary Information for "Arp2/3 complex-dependent actin remodeling is required for efficient RSV uncoating in A549 cells"

### Supplemental materials

#### Table S1. Primers used in this study.

| Target Gene | Forward (5′–3′) | Reverse (5′–3′)) | Ref. |
|---|---|---|---|
| *ACTR2* PCR primers | CAGCGAATTTGTCCATCCCAC | TAAGGTGCCAAAGAGACACTCA | |
| Sequencing primer | CATGAACCCTGGAAGTAGGC | | |
| *N* | AGATCAACTTCTGTCATCCAGCAA | TTCTGCACATCATAATTAGGAGTATCAAT | [65] |
| *NS1* | CACAACAATGCCAGTGCTACAA | TTAGACCATTAGGTTGAGAGCAATGT | [66] |
| *NS2* | TTGATGAAAAACAGGCCACA | GGGAAAGTGCCATATTTTGTGT | [67] |
| *HPRT1* | GGGAGGCCATCACATTGTAG | AATCCAGCAGGTCAGCAAAG | [68] |
| *IFNL1* | GTTCAAATCTCTGTCACCAC | TTCAGCTTGAGTGACTCTTC | [69] |
| *IFNB* | TTGACATCCCTGAGGAGATTAAGC | TCCCACGTACTCCAACTTCCA | [69] |

Abbreviations: Ref., reference*;* sgRNA, single-guide RNA; *N*, RSV nucleoprotein; *NS1*, RSV nonstructural protein 1; *NS2*, RSV nonstructural protein 2; *HPRT1*, hypoxanthine phosphoribosyltransferase 1; *IFNL1*, interferon lambda 1. *IFNB*, interferon Beta

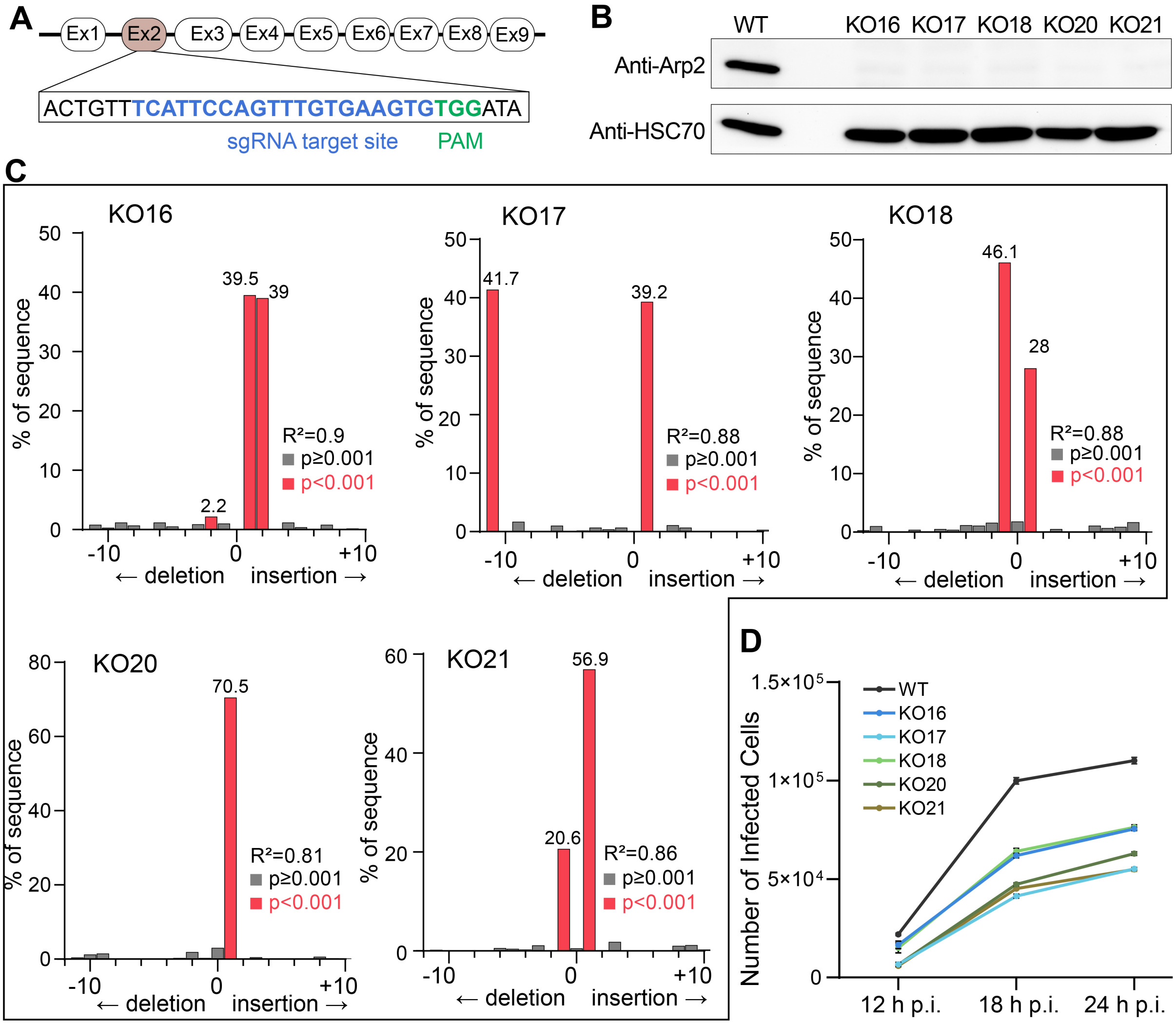


**Fig. S1 |Arp2 KO inhibits RSV infection in A549 cells at both early and late stages.**

(A) sgRNA (single-guide RNA) target site within the Arp2 (Actin-related protein 2) locus. Schematic representation of the Arp2 gene showing exons as numbered boxes (Ex1–Ex9). The protospacer (TCATTCCAGTTTGTGAAGTG) is shown in blue, and the PAM (protospacer adjacent motif; TGG) is shown in green.

(B) Immunoblot of Arp2 in A549 WT (wild-type) and Arp2 KO (knockout) cells. HSC70 (heat shock cognate 71 kDa protein) served as a loading control.

(C) TIDE sequence trace decomposition analysis of clonal cell lines derived from expanded clones, as indicated, upon A549 cell transfection with gene disruption construct PX459-Arp2. $R^2$ indicates goodness of fit.

(D) Time course of RSV-GFP infection (MOI = 3) in A549 WT and Arp2 KO cells quantified by live cell microscopy (Incucyte). Mean ± SD, n = three biological replications.

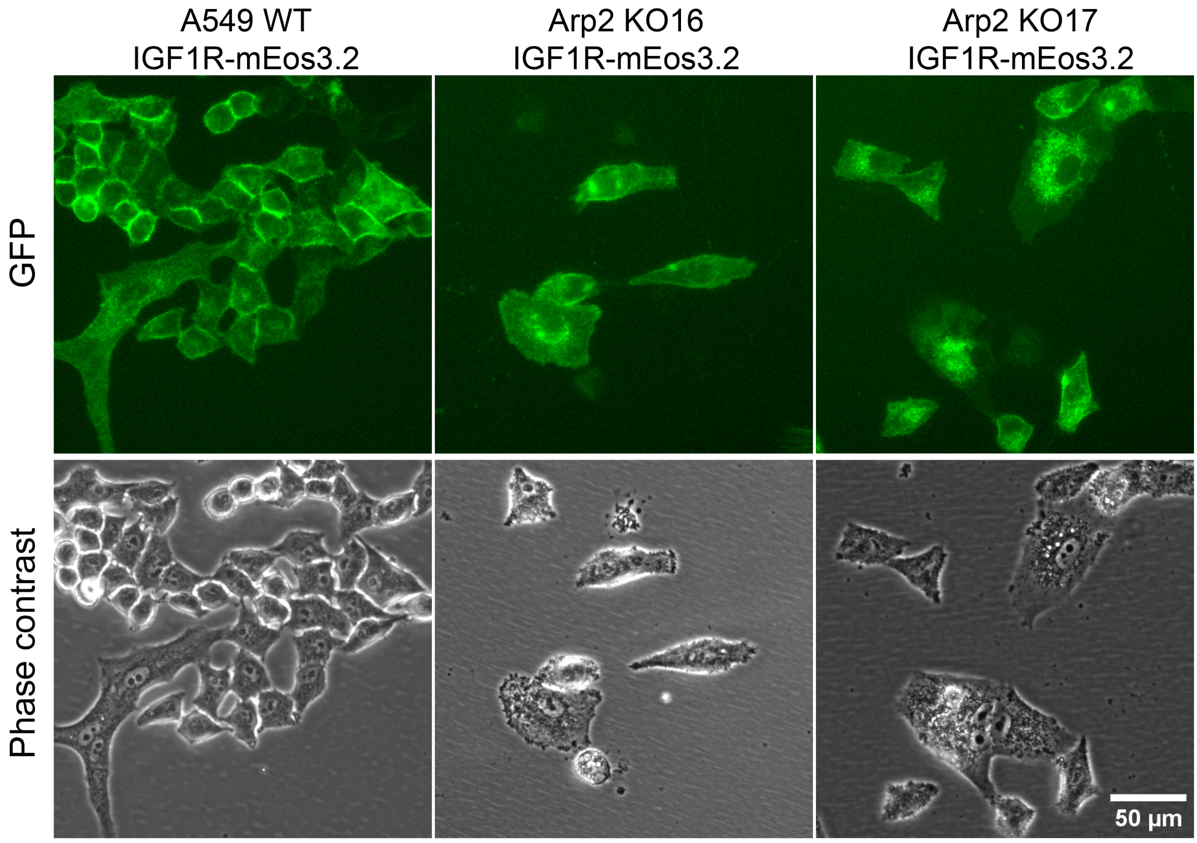


**Fig. S2 | A549 cells stably expressing IGF1R-mEos3.2.**

Representative images of WT and Arp2 KO A549 cell lines stably expressing IGF1R-mEos3.2.

Scale bar: 50 µm.

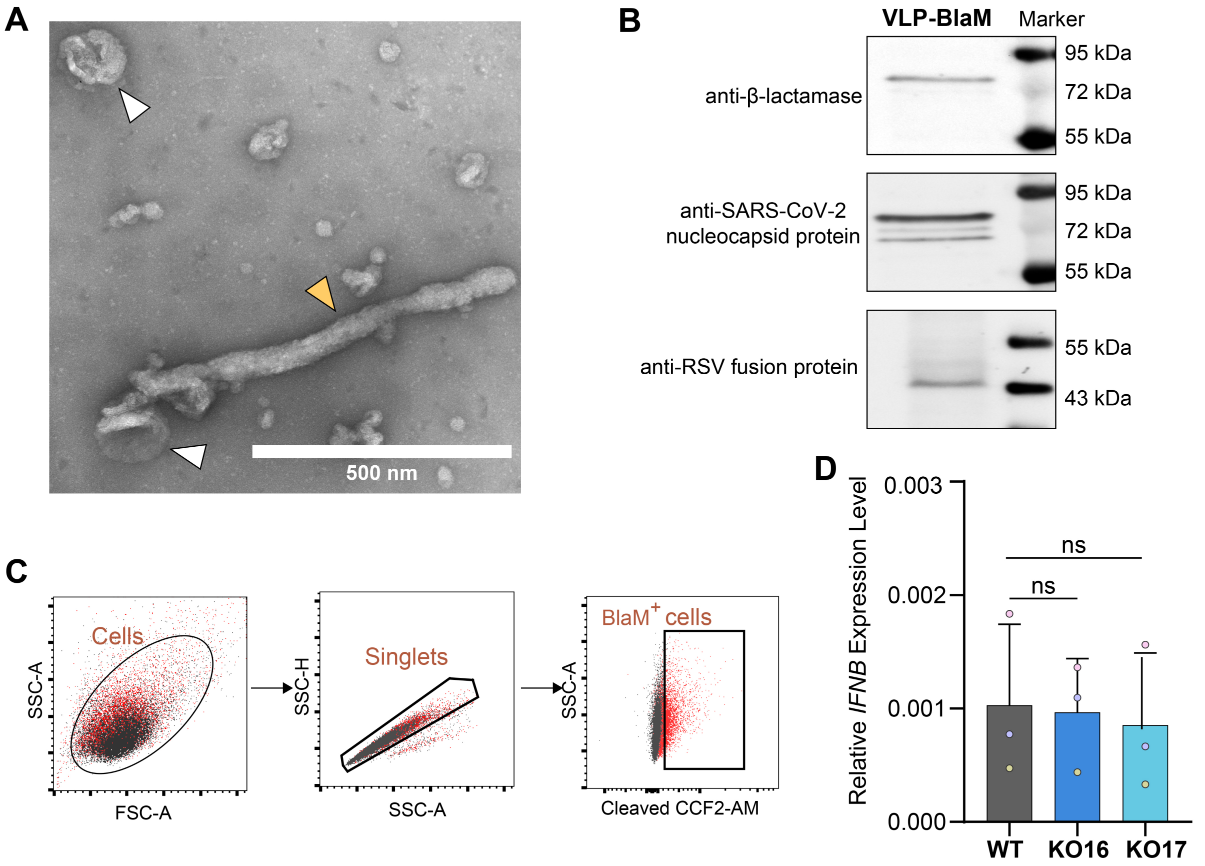


**Fig. S3 | Validation of the uncoating assay and detection of *IFNB* expression level**

(A) Representative transmission electron microscopy image of VLP-BlaM. White arrowhead indicates spherical VLP; yellow arrowhead indicates filamentous VLP.

(B) Western blot analysis of VLP-BlaM.

(C) Flow cytometry gating strategy for BlaM-positive cells. Gray dots, mock-infected controls; red dots, BlaM-VLP infected cells. FSC-A, forward scatter-area; SSC-A, side scatter–area; SSC-H, side scatter-height.

(D) RT-qPCR analysis of *IFNB* expression level at 18 h post RSV-GFP infection. The transcripts were quantified and normalized to the cellular housekeeping gene *HPRT1* (hypoxanthine phosphoribosyltransferase 1) RNA levels. Combined data from three independent experiments (three biological replicates for each experiment). Each color represents one experiment; each dot is the experiment's mean. Mean ± SD. ns, not significant (Two-way ANOVA).
